# Conserved sex-biased DNA methylation patterns target key developmental genes and non-recombining region of the guppy sex chromosome

**DOI:** 10.1101/2020.08.21.261792

**Authors:** David C.H. Metzger, Judith E. Mank

## Abstract

Understanding the regulatory mechanisms that control sexually dimorphic gene expression is a key part of understanding the processes that govern the rate of sex chromosome evolution and phenotypic sex differences. Epigenetic modifications play a major role in tissue-specific gene expression regulation and have been hypothesized to dramatically impact the speed of sex chromosome divergence. The guppy sex chromosomes are an emerging model for studying the initial stages of divergence between the X and Y. In this study, we use comparative epigenomics to identify conserved sex-specific DNA methylation patterns in gonad and muscle tissue between the Trinidadian guppy (*Poecilia reticulata*) and its sister species, Endler’s guppy (*Poecilia wingei*). We find that the oldest part of the guppy sex chromosome shows a conserved pattern of male hypomethylation, consistent with a key role in testis-specific gene expression. This pattern provides a potential mechanism for theoretical predictions that sex chromosome divergence can occur remarkably quickly in evolutionary time, and without widespread degradation of the Y chromosome gene content. Our cross-species comparative epigenomic analysis also provides a robust comparative framework to understand constraints of epigenetic programming. We observe conserved, testis-specific hypomethylated regions near key autosomal developmental genes and a potentially imprinted locus. These observations are consistent with DNA methylation in testis from other vertebrates, and suggest broad conservation of DNA methylation patterns in these regions. Our comparative framework reveals conserved DNA methylation differences between males and females across two related species, providing novel insights into the relationship between epigenetic and evolutionary processes.

## Introduction

Gene regulation is a key evolutionary process underlying phenotypic diversity. Epigenetic marks, such as DNA methylation, are dynamic biochemical mechanisms that directly influence gene expression activity. Genomic DNA methylation patterns are both environmentally pliable and heritable, thus variation in DNA methylation patterns has the potential to facilitate the rapid evolution of phenotypic diversity via flexibility in gene pathways, allowing a single genome to express different phenotypes in different environments (Jaenisch & Bird 2003; Jones 2012). Hypomethylated regions (region with low levels of DNA methylation) are typically associated with active gene expression, while hypermethylated regions (regions with high levels of DNA methylation) are associated with repressed gene expression. Consistent with a role in regulating gene expression, the dynamics of epigenetic conservation are linked with the dynamics of regulatory element turnover and chromatin structure (Zhou et al. 2017; Lowdon et al. 2016; Long et al. 2013). However, much is still unknown about the evolution and conservation of epigenetic patterns, including how epigenetic patterns evolve, and the effect of epigenetics on the rate of sequence evolution (Lowdon et al. 2016).

Epigenetic patterns can vary based on ecology, development, and experience, presenting a major challenge to identify epigenetic variation that is evolutionarily conserved (Heard & Martienssen 2014). For example, in higher vertebrates, parental epigenetic marks are reprogrammed in the germline and early stages of embryogenesis, thus stable epigenetic variation may be quite rare (Ortega-Recalde & Hore 2019) In contrast, transgenerational epigenetic variation might be more prominent in organisms that lack epigenetic reprogramming. For example, genomic DNA methylation patterns of zebrafish sperm are maintained after fertilization, and are thought to act as a template during early gametogenesis (Potok et al. 2013; Jiang et al. 2013; Skvortsova et al. 2019; Ortega-Recalde et al. 2019). However, it is unknown how common the preservation of paternal DNA methylation patterns is among teleosts, as medaka undergo extensive reprogramming of DNA methylation, similar to mammalian systems (Wang & Bhandari 2019).

Multi-species comparisons of tissue-specific DNA methylation patterns are a powerful way of overcoming the challenges described above to identify conserved epigenetic patterns of functional regulatory elements that underlie gene expression variation (Zhou et al. 2017; Lowdon et al. 2016; Long et al. 2013). This approach is particularly effective when coupled with comparisons of discrete phenotypic polymorphisms, such as sexual dimorphism. In this study, we assess sexually dimorphic patterns of DNA from two closely related species of Poeciliids, the Trinidadian guppy (*Poecilia reticulata*) and its sister species, Endler’s guppy (*Poecilia wingei*) (Pollux et al. 2014). Sexual dimorphism is a striking example of intraspecific variation of complex phenotypes, and both *P. reticulata* and *P. wingei* exhibit pronounced sex differences in size, color and behavior (Poeser et al. 2005). Sexually dimorphic traits are the result of gene expression differences between males and females (Mank 2017). Moreover, although many sexually dimorphic traits are encoded by autosomal genes, genes located on sex chromosomes often play a disproportionately large role in regulating sexually dimorphic traits (Mank 2017). It is therefore perhaps not surprising that the sex chromosomes are often a hotspot of sex-specific methylation (Metzger & Schulte 2018; Liu et al. 2010). However, although the link between sex-specific methylation and highly diverged heteromorphic sex chromosomes is well established, we know far less about sex-specific epigenetic regulation of less diverged, homomorphic sex chromosomes.

*P. reticulata* and *P. wingei* share the same XY sex chromosome system on Chromosome 12 which is thought to have originated at least 20Mya (Darolti et al. 2019; Rabosky et al. 2018; Tripathi, Hoffmann, Willing, et al. 2009). While a sex determination gene has not yet been identified, linkage maps have localized the sex determination locus (SDL) to be at the distal end of Chromsome 12 (Tripathi, Hoffmann, Willing, et al. 2009). Stratum I (20-26Mb) is the oldest, non-recombing region of this chromosome pair exhibiting the most divergence between the X and Y and is ancestral to *P. reticulata* and *P. wingei*. (Darolti et al. 2019, 2020; Almeida et al. 2020). A more recent region of recombination suppression (Stratum II) has also been identified proximal to Stratum I. The size of Stratum II differs between *P. reticulata* and *P. wingei* and among wild *P. reticulata* populations (Darolti et al. 2019; Wright et al. 2017; Almeida et al. 2020), and appears to have formed independently numerous times. Moreover, there is little evidence of gene loss from the Y chromosome, either in Stratum I or the remainder of the sex chromosome (Darolti et al. 2020). Therefore, it is possible that sex-biased gene expression patterns between males and females in these species may be due to sex-biased epigenetic variation instead of genetic variation between sex chromosomes. The shared homomorphic sex chromosome system and pronounced dimorphism in these species make *P. reticulata* and *P. wingei* an ideal comparative system to assess both the evolutionarily conserved patterns of sexually dimorphic DNA methylation patterns, and the relationship between DNA methylation and sex chromosome divergence.

In this study we use a comparative epigenomics approach to identify conserved regions of *P. reticulata* and *P. wingei* tissue-specific DNA methylation patterns in gonad and muscle tissue from both sexes. We find that testis has a highly specialized genomic DNA methylation pattern compared to ovary and muscle, which exhibit relatively little sex-specific DNA methylation. Testis-specific DNA methylation patterns are typified by distinct clustering of hypomethylated loci in regions that are consistent with a regulatory role in activating critical developmental genes and suggest a constraint on the epigenetic programming in these regions. We also discover testis-specific hypomethylated near the boundaries of Stratum I on the sex chromosome. These regions are consistent with the ancestral regions of recombination suppression near the SDL and suggests that the conserved hypomethylated regions on the sex chromosome may be involved in regulating sex-biased gene expression underlying sexually dimorphic phenotypes. Moreover, these data suggest a potential mechanism for theoretical predictions (Rice 1984) that sex-limited regulatory features can be established remarkably quickly in evolutionary time, and without widespread degradation of genes on the Y chromosome.

## Results

Using whole genome bisulfite sequencing (WGBS) data from male and female gonad and muscle tissue from two Poeciliid sister species, *P. reticulata* and *P. wingei*, we observed that the testis has a higher average DNA methylation level (~72%) compared to the ovary and muscle tissue in both sexes, which exhibit similar genome wide levels of DNA methylation (~68%) (fig. 1A). A principle components analysis revealed the testis contributed the most variance in DNA methylation patterns (31%, PC1), suggesting a highly specialized DNA methylation pattern in the testis compared to other tissues and that DNA methylation patterns in testis are conserved between these two species (fig. 1B & C). Species-specific DNA methylation patterns were the second largest factor contributing to the variance (28%, PC2) in DNA methylation patterns (fig. 1B). Differences in tissue-specific patterns between ovary and muscle tissue accounted for 4% (PC3) of the variance (fig. 1C).

**Figure 1:**
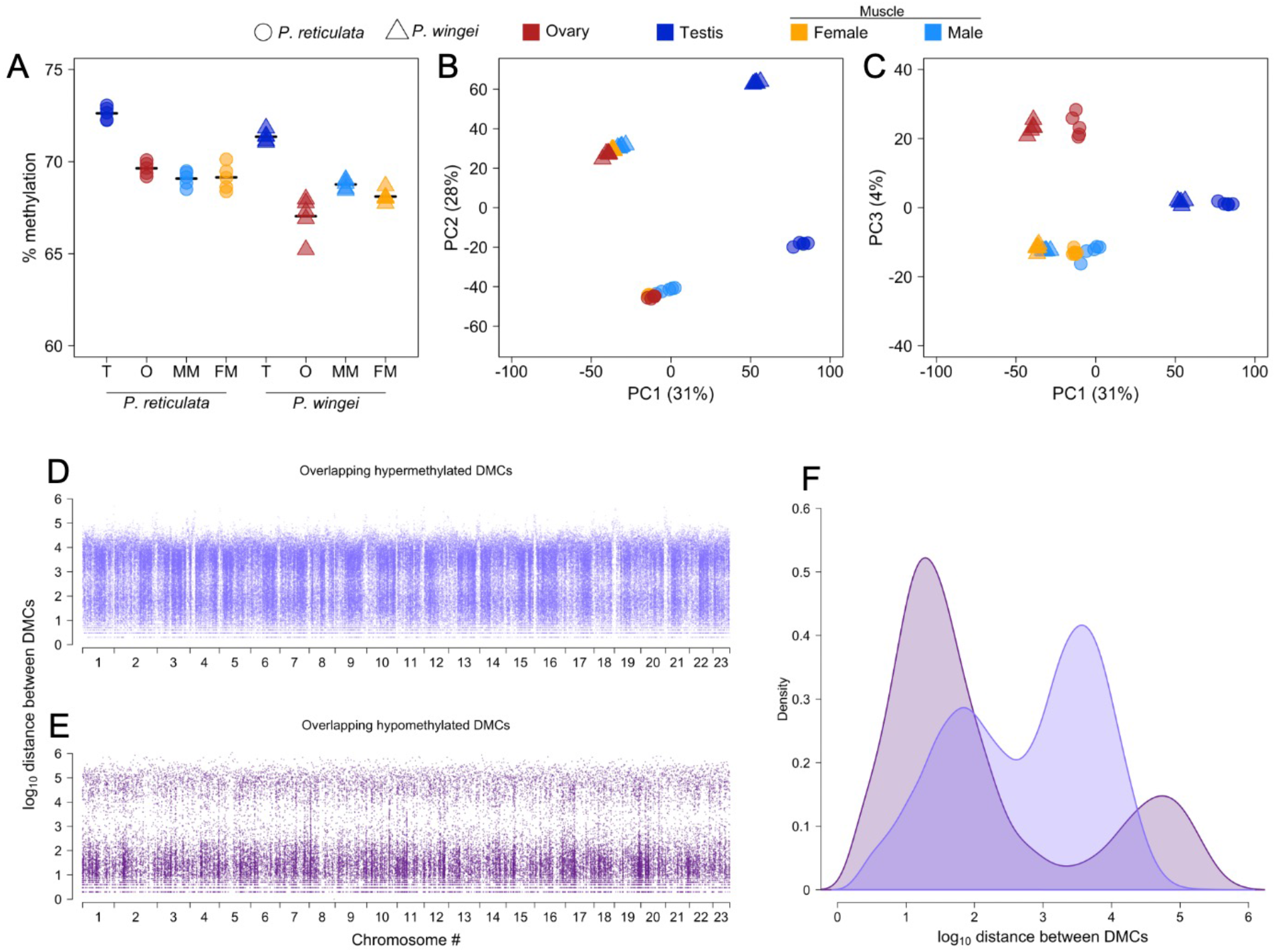
Conservation of testis-specific DNA methylation patterns between *Poecilia reticulata* and *Poecilia wingei*. (A) Average genome-wide DNA methylation levels for testis (T), ovary (O), male muscle (MM), and female muscle (FM) tissue from *P. reticulata* (circles) and *P. wingei* (triangles). Each point is the average genome-wide DNA methylation for each sample. The horizontal line is the mean DNA methylation level for the five samples in each category. (B, C) Principal component analyses (B: PC 1 and PC 2, C: PC 1 and PC 3) of DNA methylation from whole genome bisulfite sequencing. Colors and shapes designating sample type are the same as in panel A. (D-E) log_10_ genomic distance between hypomethylated (D) or hypermethylated (E) differentially methylated CpG loci that are conserved between *P. reticulata* and *P. wingei*. (F) Density distribution of the log_10_ distance between DMCs; hypomethylated (purple), hypermethylated (blue).

One advantage of using WGBS is that it allows for the analysis of site-specific information about DNA methylation levels of individual CpG loci. In general, ~20% of the CpG sites were differentially methylated between testis and other tissues (table 1). In order to differentiate between unmethylated and methylated cytosines, bisulfite sequencing relies on the biochemical mutation of all unmethylated cytosines to thymines, while methylated cytosines are unaffected. When mapping bisulfite treated sequences to a reference genome, it can be difficult to distinguish bisulfite converted mutations from genetic variation between the sample and the reference genome. However, similar to the PCA analysis, we identified very few significantly differentially methylated cytosines (DMCs) between male and female muscle tissue, which strongly indicates that the effects of genetic variation between the sample DNA and the reference genome is minimal (table 1, fig. S3-7). These results suggest the testis has a unique DNA methylation profile, and this finding is consistent with the results from the PCA analysis. We therefore focused our subsequent analyses on the unique DNA methylation patterns of the testis, specifically the comparison of testis and ovary, the latter having the most CpG cites that met the minimum criteria for the pairwise comparison analysis (table 1).

**Table 1:** Summary of DMCs from pairwise comparisons of tissue and sex in two closely related species.

|  | *P. reticulata* |  | *P. wingei* |  | Overlapping sites |  |
| --- | --- | --- | --- | --- | --- | --- |
|  | # CpGs | # DMCs | # CpGs | # DMCs | #CpGs | #DMCs |
| Testis v ovary | 3,688,554 | 764,125 (20.7%) | 2,485,952 | 564,732 (22.7%) | 1,563,847 | 231,015 (15%) |
| Testis v muscle | 1,681,470 | 319,799 (19%) | 2,320,394 | 424,669 (18%) | 798,688 | 95,903 (12%) |
| Ovary v muscle | 1,989,533 | 101,716 (5%) | 1,644,089 | 74,211 (4.5%) | 727,698 | 12,146 (1.7%) |
| male v female muscle | 1,024,996 | 2,915 (0.3%) | 1,549,105 | 3,288 (0.2%) | 430,771 | 18 (0.002%) |

Differential methylation patterns between testis and ovary are highly correlated between *P. reticulata* and *P. wingei* (R^2^ = 0.617, p < 0.001, fig. S1). Previous studies comparing DNA methylation patterns between species have shown that conserved genomic regions of DNA methylation between species represent shared key regulatory elements involved in differentiation (Zhou et al. 2017). To identify putative regulatory elements that could be involved in regulating sexually dimorphic phenotypes, we focus our analysis on DMCs that are conserved between *P. reticulata* and *P. wingei*.

### Methylation clusters

To identify discrete clusters of DMCs that could represent DNA methylation dependent regulatory regions, we searched for clusters of hypo- and hypermethylated DMCs by calculating the distance between neighboring DMCs and analyzed the spatial distribution of clusters in the genome. If hypo- and hypermethylated DMCs are evenly dispersed throughout the genome, hypermethylated sites would be expected to cluster more tightly due to greater abundance. However, we observed the opposite pattern. The majority of hypomethylated DMCs occur in tight clusters, while the organization of hypermethylated DMCs was less structured and more dispersed (fig. 1D-E, fig. S2).

We used a two-factor approach to identify discrete genomic windows containing clusters of hyper or hypomethylated DMCs. First, we normalized the total number of DMCs in a window to the total number of CpG sites that were sequenced in that window and that passed the minimum quality thresholds described above. However, because CpG loci are not evenly distributed throughout the genome, the proportion of DMCs can be skewed by the total number of CpG sites in that window. Therefore, we also used a binomial test to identify regions with a greater abundance of DMCs. We then focused on regions where both the normalized DMC proportional value and the FDR corrected q-value from the binomial test exceed their respective 95% confidence interval. Using this conservative approach, we identified 23 outlier genomic regions containing both higher proportions and greater numbers and of hypomethylated DMCs compared to other regions of genome (fig. 2A, table S2). While 43 windows of hypermethylated sites were also found (fig. 2A, table S3), these windows were not among the top candidate regions.

**Figure 2:**
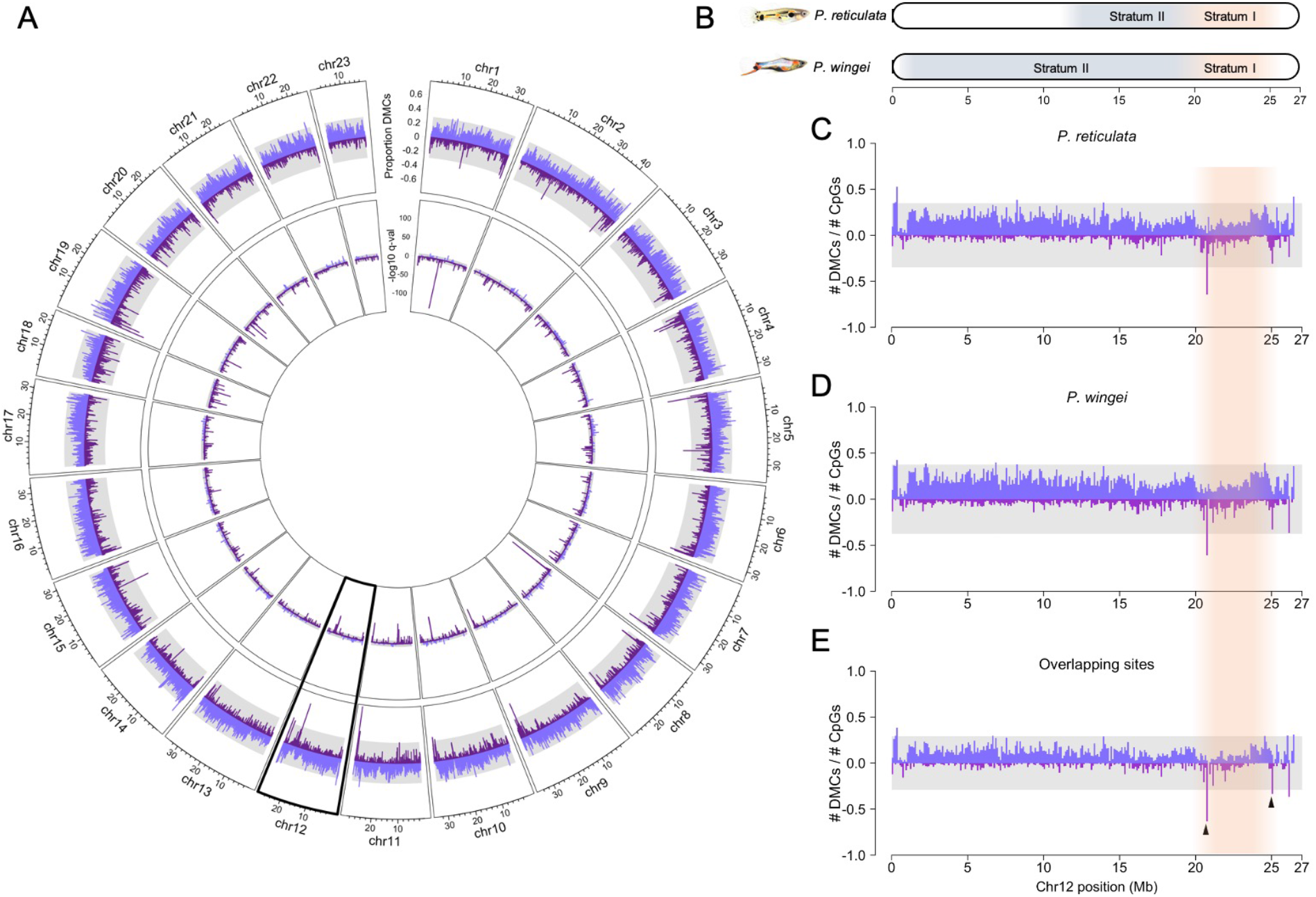
Genomic distribution of differentially methylated CpG loci (DMCs). (A) Circos plot of all 23 guppy chromosomes. The outer ring depicts the proportion of DMCs between testis and ovary within a 100kb window that are conserved between *P. reticulata* and *P. wingei*: hypermethylated (blue), hypomethylated (purple). The inner circle is the log_10_ FDR corrected q-value of a binomial test on the number of hyper- or hypomethylated DMCs between testis and ovary in 100 kb windows. (B) Schematic of the guppy sex chromosome (Chromosome 12) depicting Stratum I (orange) and Stratum II (blue). Scale bar represents chromosome position in mega base pairs. (C-E) Proportion of hypermethylated and hypomethylated DMCs between testis and ovary in 100kb windows of chromosome 12 for *P. reticulata* (C), *P. wingei* (D), and DMCs that overlap between the two species (E). Shaded grey areas represent the 95% confidence interval. Shaded orange area represents the location of Stratum I, and the black arrows in panel E highlight conserved regions of enriched hypomethylation near the boundaries of Stratum I. Negative values depict hypomethylated DMCs for visualization purposes only.

Conserved DNA methylation patterns are often located in the CpG islands of promoter regions and other regulatory elements. In mammals, many CpG islands (CGI) are located near transcription start sites and there is a high correlation between hypomethylated regions and predicted CGIs. In contrast, CGI predictions in poikilothermic vertebrates often reside away from gene promoters and are poor predictors are hypomethylated regions (Long et al. 2013). This is likely due in part to the fact that algorithms to detect CGIs are based on the genomic features (eg. CpG density, G+C content, genome assembly quality) of mammals, which are not conserved across all vertebrates (Long et al. 2013). Thus, we use the term non-methylated island (NMIs) (Long et al. 2013; Lowdon et al. 2016) to describe conserved regions of hypomethylated CpG clusters, which are conserved features of orthologous promoters and distal regulatory elements. Many of the NMIs are associated with genes involved in key developmental processes typically associated with hypomethylated valleys (e.g. *hox* gene cluster on chr8), nervous system development (e.g. ephrin type-A receptor 7-like on chr11 and *fgf11a* on chr18), or that are orthologous to genes that have been previously described as components of imprinted clusters in mammalian systems (e.g. ENSPREG00000000764 on chr1).

We also identified regions with multiple CpG methylated loci in discrete regions, or differentially methylated regions (DMRs). The majority of DMRs were hypomethylated in testis compared to ovary. We identified 2,480 hypomethylated and 759 hypermethylated DMRs in *P. reticulata*, 1,709 hypomethylated and 363 hypermethylated DMRs in *P. wingei*, and 867 hypomethylated and 93 hypermethylated DMRs that were present in both species. These results are consistent with our previous analyses, as more clustering of hypomethylated DMCs should result in more hypomethylated DMRs.

We performed a GO enrichment analysis for genes within 5kb of a DMR, focusing on processes associated with genes near hyper or hypomethylated DMRs separately. 24 GO terms were significantly enriched for genes located near hypomethylated DMRs. Many of these GO terms were associated with cell adhesion, signal transduction, and tissue development processes. In contrast, protein phosphorylation was the only significantly enriched GO term for genes located near hypermethylated DMRs (tables S4 & S5).

### NMIs on the sex chromosome

Analysis of DMCs on Chromosome 12 identified a testis-specific region of hypomethylation at the boundary of Stratum I and Stratum II of the guppy sex chromosome (fig. 2B-E). Stratum I (21-26Mb) is ancestral to *P. reticulata* and *P. wingei* and although there is no evidence of gene loss from the Y chromosome, there is significant divergence between the X and Y. The sex determining locus, although not yet known, has been mapped to the distal end of Chromosome 12 in Stratum I, and the hypomethylated windows on Chromosome 12 are particularly interesting because they are located at the boundaries of this region in both *P. reticulata* and *P. wingei* (fig. 2C & D). We have previously noted that the boundaries of Stratum I show the greatest pattern of X-Y divergence and accumulation of male-specific sequence, consistent with being the oldest region of suppressed recombination between the X and Y chromosomes (Almeida et al. 2020; Darolti et al. 2020). Stratum II is younger, formed independently in each species, and varies in size between species and among *P. reticulata* populations (fig. 2B) (Wright et al. 2017; Darolti et al. 2019, 2020; Almeida et al. 2020). In contrast to the hypomethylated patterns observed in Stratum I, there are relatively few hypomethylated regions in Stratum II, and the distribution differs between species (fig. 3B-D). This observation is consistent with Stratum I being the older evolutionary stratum, and that X and Y divergence in this region is ancestral to these species. It is important to note that we do not find these differential methylation patterns in muscle tissue, and that the number of DMCs on each chromosome was correlated with the chromosome length, indicating there was no apparent enrichment of DMCs at the chromosome level (fig. S8), and that X and Y chromosome divergence was not biasing sex-specific DNA methylation patterns to the sex chromosome.

The hypomethylated window between 20.7 and 20.8 Mb of Chromosome 12 and near the boundary of Stratum I contains an NMI downstream of the NLR family CARD domain-containing protein 3-like (*nlrc3-like*) gene, a negative regulator of the Toll-like receptor pathways and innate immune response (fig. S9) (Schneider et al. 2012). Although it is possible that this region could function as a regulatory element for the expression of *nlrc3-like*, methylation sensitive regulatory elements are typically located upstream near promoter regions of genes. To identify other sequence features associated with the hypomethylation clusters near the sex determining locus, we used a BLAST search of the sequence 5kb upstream and downstream of the DMCs. Using this approach, we identified a region of ~1.7kb with sequence homology to genomic regions in other teleost species but with no known annotation. Using the repeat masking function in *repbase*, we identified this sequence as a potential non-LTR retrotransposon rex1.

To determine whether DMRs could have a functional role in regulating sex-linked genes, we compared a list of 42 recently identified sex-linked genes on the guppy sex chromosome (Darolti et al. 2020) with genes located within 5kb of a DMR that was present in both species. In total, we identified 73 genes near hypermethylated DMRs and 658 genes near hypomethylated DMRs. Six of the 42 sex linked genes (*dnajc25, skiv212, arhgap24, nup214, npr2*, and *tsc1*) were within 5kb of a DMR on Chromosome 12. Interestingly, the DMR with the highest test statistic on Chromosome 12 was near *dnajc25* in Stratum I, which is one of five candidate genes with a Y-specific gametolog (Potok et al. 2013; Jiang et al. 2013).

## Discussion

In this study, we use a comparative epigenomic approach to identify conserved, sex-specific DNA methylation patterns in gonad and muscle tissue from two sister species of Poeciliids, *P. reticulata* and *P. wingei*. Consistent with other vertebrates, we observe conserved DNA methylation patterns in genomic regions containing key developmental processes, suggesting the paternal genome is important for establishing DNA methylation patterns of critical developmental genes in the embryo. We also identify a conserved region of testis-specific DNA methylation consistent with a role in regulating sex-specific regulatory processes. Genomic regions containing conserved DNA methylation often contain gene regulatory elements that control gene expression (Hernando-Herraez et al. 2015), thus the conserved DNA methylation patterns identified in this study may play a role in regulating sexually dimorphic gene expression patterns.

### Conserved hypomethylation of developmental genes in testis

Comparison of genomic DNA methylation levels between male and female gonad and muscle tissue revealed a higher genome wide DNA methylation level in testis compared to ovary or muscle, which has been observed in other vertebrates (Potok et al. 2013; Jiang et al. 2013). We also identified more DMCs in testis compared to other tissues, suggesting that the genomic DNA methylation pattern of testis is highly specialized compared to muscle and ovary.

Despite observing higher levels of DNA methylation in testis, we found more genomic regions enriched for hypomethylated clusters in testis. The clustering of DNA hypomethylation in testis is consistent with these regions being associated with positive regulation of testis-specific gene expression. In other vertebrates, conserved hypomethylated regions are also located near regulatory elements with key roles in tissue-specification (Tena et al. 2014). During spermatogenesis some histones are replaced by protamines, the remaining histones have been associated with hypomethylated promoters of key developmental genes (Champroux et al. 2018). In this study, we used two separate approaches (clustering and dmrseq) to identify genomic regions containing clusters and both approaches identified genes related to developmental processes. Taken together, our results are consistent with the observation that paternal epigenetic patterns are important regulators of key developmental genes (Potok et al. 2013; Jiang et al. 2013; Hammoud et al. 2009), and that NMI’s near developmentally important genes in the male gonad are conserved among vertebrates. This suggests a constraint on epigenetic patterns of genes necessary for proper vertebrate development (Tena et al. 2014)

The most significantly enriched region for hypomethylated loci is on Chromosome 1 and contains an NMI on a gene (ENSPREG00000000764) orthologous to *rtl1* (also known as *peg11*) in *Xiphophorus maculatus* and *rtl2* (also known as *peg10*) in mammals (fig. S10). Both *rtl1* and *rlt2* are components of maternally imprinted gene clusters essential for proper implantation and development in mammals (Ono et al. 2006; Luo et al. 2016; Yu et al. 2018), and are thought to have been important for the evolution of the placenta (Yu et al. 2018; Sekita et al. 2008). Genomic imprinting is the parent of origin-specific regulation of gene expression achieved through allele-specific DNA methylation patterns. The DNA methylation of ENSPREG00000000764 that we observe in testis is ~50% and is hypomethylated compared to ovary (fig. S10), which is consistent with the expectation of a maternally imprinted gene. Although fish lack the *dnmt3l* gene (Yokomine et al. 2006), necessary for establishing DNA methylation status of imprinted loci in mammals (Hata et al. 2002), there is some limited evidence for genomic imprinting in livebearers (Liu et al. 2010; Saldivar Lemus et al. 2017). Moreover, livebearers are a classic model for studying the evolution of the placenta, and the conserved NMI of ENSPREG00000000764 suggests that it could be an important component evolution and development of the placenta (Pollux et al. 2014). While these data suggest that imprinting may indeed exist in some live-bearing fish, whether or not this conserved region of hypomethylation is a novel imprinted locus requires further investigation.

### Conserved testis-specific hypomethylation near sex determining locus

In vertebrates, sexually dimorphic DNA methylation patterns have been shown to be involved in environmentally sensitive sex determination mechanisms (Chen et al. 2014; Shao et al. 2014; Navarro-Martín et al. 2011; Anastasiadi et al. 2018), X-chromosome inactivation (Liu et al. 2010), and genomic imprinting (Barlow & Bartolomei 2014). Much of what is known about sex-biased DNA methylation patterns comes from species with degenerated heteromorphic sex chromosomes where sex-biased DNA methylation patterns are enriched on the sex chromosome (Metzger & Schulte 2018; Liu et al. 2010). In mammalian systems, enrichment of sex-biased DNA methylation patterns on the sex chromosome may be largely due to dosage compensation, and the role of epigenetic processes in X-chromosome inactivation (Liu et al. 2010). However, enrichment of sex-biased DNA methylation on sex chromosomes has also been observed in the absence of dosage compensation despite extensive Y degeneration (Metzger & Schulte 2018). In guppies, the sex determining locus (SDL) is located near the distal end of chromosome 12 and is thought to have originated ~20Mya (Darolti et al. 2019; Tripathi, Hoffmann, Weigel, et al. 2009). Although genetic divergence has accumulated on the Y chromosome around the region containing the SDL in Stratum I, there is little evidence of gene loss on the Y (Wright et al. 2017; Darolti et al. 2019; Almeida et al. 2020).

Although we did not observe enrichment of sex-biased DNA methylation of the guppy sex chromosome relative to the autosomes, we did identify a conserved hypomethylated region near the boundary of Stratum I and Stratum II (Darolti et al. 2020), between 20.7 and 20.8 Mb of Chromosome 12 and two regions at the distal end of Stratum I near the boundary with the distal pseudoautosomal region at 26 Mb (fig. 2E). These regions have recently been highlighted as the oldest non-recombining regions of the sex chromosome (Almeida et al. 2020; Darolti et al. 2020). The testis-specific hypomethylation in these regions are present in both *P. reticulata* and *P. wingei* and are more prominent when examining only the conserved DMCs between the two species (fig. 2C-E) suggesting that the DNA methylation profile of this region predates the divergence of these two species and may coincide with recombination suppression between the X and Y.

Variation in DNA methylation patterns and histone modifications are associated with chromatin structure and recombination landscapes (Mirouze et al. 2012), which has been hypothesized to play an important role in initial recombination suppression between emerging sex chromosome pairs (Gorelick 2003). This hypothesis posits that hypermethylation of a novel SDL establishes sex biased gene expression, simultaneously initiating heterochromatinization and recombination suppression in this region, which gradually expands to form larger strata (Gorelick 2003). While guppies are thought to have expanding strata, we found the opposite pattern than would be predicted if recombination suppression was being driven by DNA methylation. While we do observe global hypermethylation in testis, this pattern is neither localized to strata or unique to the sex chromosome and is therefore unlikely to have been the direct cause of recombination suppression of Stratum I. In addition, we see relatively little conserved hypomethylation of Stratum II. Stratum II has evolved independently between *P. wingei* and *P. reticulata* (Darolti et al. 2019), and between different populations of *P. reticulata* (Almeida et al. 2020). The absence of conserved hypomethylation similar to what we found in Stratum I suggests that conserved hypomethylation is not a direct cause or consequence of recombination suppression between sex chromosomes in these species.

### DNA methylation of transposable elements on the sex chromosome

At least one hypomethylated region on the sex chromosome may be associated with the non-LTR retrotransposon *rex1*. Transposable elements (TEs) are selfish genetic sequences that can replicate and move to different genomic regions and can modify gene function by providing new or disrupting existing regulatory or coding sequences of genes (Volff 2006). Due to the potential destructive nature of TEs, some studies suggest that DNA methylation originated as a genome defense mechanism to suppress transposable element activity and was later adapted for other purposes such as genomic imprinting (Deniz et al. 2019). Thus, the movement and activity of TEs can also have localized effects on epigenetic patterns that can affect expression of nearby genes (Hollister & Gaut 2009).

Because of their ability to move genetic material around the genome and alter expression patterns, TEs are thought to be a potential mechanism to promote turnover of sex determining loci while simultaneously suppressing recombination (Furman et al. 2020). Consistent with this hypothesis, TEs have been found at the boundaries of nonrecombining regions of recently established regions of suppressed recombination in mammals and birds (Iwase et al. 2003; Xu et al. 2019), and are also enriched in Stratum I in guppies (Almeida et al. 2020). Therefore, it is not surprising to find DNA methylation associated with a putative TE in the non-recombining region of the sex chromosome, however it is unclear why there is testis-specific hypomethylation of this TE which is consistent with an activation of the TE. One possibility is that this hypomethylation cluster represents a novel regulatory element that was brought to this region by the rex-1 retrotransposon and has since acquired male-biased function that is activated in testis and turned off in other tissues. It is also possible that male-biased activation of TEs could lead to further replication and expansion of the non-recombing region of the guppy sex chromosome, a process that may be occurring in *P. wingei*.

### Conclusion

Epigenetic variation has the potential to play a key role in modulating phenotypic diversity though differential regulation of gene expression patterns. However, identifying conserved epigenetic variation can be challenging due to the reprogramming of epigenetic patterns each generation. Using a comparative epigenomic approach, this study provides novel insights into the relationship between epigenetic and evolutionary processes, and the conservation of epigenetic programming. Most striking was the conservation of testis-specific hypomethylation regions in Stratum I of the guppy sex chromosome. These data suggest a potential mechanism for the rapid, sex-limited divergence in this region without extensive Y chromosome degradation. Conserved hypomethylated regions are also associated with key developmental genes, a pattern consistent with other vertebrates suggesting a constraint on the evolution of developmentally important DNA methylation patterns. Given these constraints, the conserved hypomethylated regions in Stratum I of the guppy sex chromosome is consistent with a key role in regulating the sex-biased expression in this region. Taken together, these data demonstrate how using a cross-species comparative epigenomics approach can reveal novel candidate epigenetic regulatory features and provide insight on the dynamics of epigenetic programming and genetic divergence.

## Materials and Methods

### Sample collection and sequencing

We collected adult male and virgin females from laboratory populations of *Poecilia reticulata* and *Poecilia wingei*, maintained at the University College of London aquatics facility under institutional licence (PEL PCD 70/2716). The *P. reticulata* lab strain is an outbred laboratory population originating from a high predation population in the Quare River, Trinidad (Kotrschal et al. 2013). The *P. wingei* laboratory population was obtained from a UK fish hobbyist. All fish were maintained under a standard diet of fancy guppy flake and artemia, at 25 °C and a 12h-12h light cycle. Following a lethal dose of MS-222, we immediately dissected muscle and gonad tissue which were then stored in 95% ethanol. Muscle tissue was taken from the anal pore to the base of the tail with the skin and bone tissue removed.

Inter-individual DNA methylation variation can be attributed to genetic differences as well as environmental factors and life experiences (Jaenisch & Bird 2003). To minimize the effects of inter-individual variation, we pooled tissue from five individuals for each sequencing library. Five libraries were created for each tissue for both species, resulting in a total of 25 male and 25 female fish sampled for each species. The same individuals that were pooled for the gonad samples were also pooled for muscle. We stored tissue at 4 °C for 24-72 hours prior to extracting DNA with a DNAeasy Blood and Tissue Kit (Qiagen) with on-column RNase A treatment following the manufacturer’s protocol. Resulting genomic DNA samples were frozen at −20 °C, and then shipped overnight on dry ice to the McGill University and Génome Québec Innovation Centre for bisulfite conversion, library preparation, and sequencing on the Illumina HiSeqX PE150 platform.

Quality filtering and adapter trimming was performed using Trimmomatic v0.38 (Bolger et al. 2014). Reads were scanned with a 4-base sliding window and discarded when the average phred score was < 15. Leading/trailing bases with a phred score < 3 were also removed. Data quality was assessed using FastQC v0.11.7. Approximately 75 million reads were retained for each library after quality filtering. The *P. reticulata* reference genome was obtained from NCBI Genome database (Guppy_female_1.0+MT, assembly accession GCA_000633615.2), correcting for the large inversion on Chromosome 12 (Almeida et al. 2020; Darolti et al. 2020). Trimmed and filtered reads were mapped to the reference genome using Bismark v0.20.0 (Krueger & Andrews 2011) and the BAM alignment files were sorted using samtools v1.9 (Li et al. 2009). The average mapping efficiency was ~67% and covered ~2 million CpG sites (~16.8% of the total number of CpG sites in the *P. reticulata* genome) at a depth of at least ten reads.

### DNA methylation analysis

DNA methylation analysis was conducted using R v3.5.3 (R Core Team 2019) and the R package *methylKit* v1.8.1 (Korthauer et al. 2019). Methylation percentages were calculated using the*process Bismark Aln* function. For the analysis of individual CpG loci, sites with < 10 reads or that were in the 99.9th percentile of coverage were removed from the analysis. Principle component analysis was perform using only sites that passed the minimum quality threshold in all samples using the *prcomp* function. To perform pairwise comparisons between tissues, a logistic regression model was used with the *calculateDiffMeth* function with a correction for over dispersion. A Chi-square test was used to identify significantly differentially methylated cytosines (DMCs) between tissues. Sequencing lane was included as a covariate. False discovery rate (FDR) corrected p-values (q-values) were calculated using the sliding linear model method (SLIM) with a maximum q-value threshold of 0.05 and a minimum change in percent methylation of 20%. We focused primarily on comparing DNA methylation patterns between testis and ovary because this comparison had the greatest number of CpG sites that met these criteria. Data for the other tissues can be found in the supplemental material. Pairwise comparisons were performed to identify individual CpG loci with sex-specific DNA methylation patterns.

### Clusters of differentially methylated CpG loci

In order to identify clusters of hypomethylated and hypermethylated DMCs, we employed an approach analogous to a kataegis analysis, typically performed to identify localized clusters of genetic mutations based on the genetic distance between sites (Nik-Zainal et al. 2012). Instead of calculating the distance between mutations, we calculated log_10_ distance between neighboring DMCs to determine localized clustering of hypomethylated and hypermethylated sites. We then built rainfall plots to visualize genome-wide patterns of hypomethylated and hypermethylated DMC clustering patterns.

The number of hypomethylated and hypermethylated DMCs in a window was normalized to the total number of CpG sites retained after quality and depth filtering in that window. Because some windows contain more CpG sites, normalizing to the number of sequenced CpG sites provides an estimate to the amount of differential methylation in that region. This normalization was conducted separately for the total number of DMCs, the number of hypomethylated DMCs, and the number of hypermethylated DMCs. A 95% confidence interval was calculated using the normalized values for all DMCs (excluding Chromosome 12).

We used a sliding window approach to characterize the proportion of hypomethylated or hypermethylated CpG loci across the genome using 100 kb consecutive non-overlapping windows. A binomial test was used to determine whether the number of hypomethylated or hypermethylated sites in a given window was greater than the number that would be expected by chance given the number of CpG sites that were sequenced and passed the quality thresholds described above. Only windows that contained >10 CpG sites were included in the analysis. A 95% confidence interval for the p-value statistics resulting from the binomial test for the total number of differentially methylated sites in a window was calculated to identify outliers of either hypermethylated or hypomethylated windows.

### Differentially Methylated Regions

Although differential methylation of individual CpG loci can have significant regulatory effects, methylation sensitive regulatory elements are often controlled by the DNA methylation status of multiple CpG loci in discrete regions. To identify these differentially methylated regions (DMRs) we used *dmrseq* v1.2.5 (Korthauer et al. 2019). Unlike other programs that require the user to provide an arbitrary window size, *dmrseq* uses a more dynamic approach in which an initial scan of the methylation data is performed to identify putative DMRs. Using this approach, the regions can vary in size and the size is determined by the data instead of the user. DMRs were characterized as regions with a p-value < 0.05 and a test statistic greater than 0.2.

### GO enrichment of DMRs

Genes associated with DMRs were used for a GO enrichment analysis. To identify genes associated with DMRs, the *subsetByOverlaps* function was used to subset the GTF file for genes within 5kb of a DMR. A hypergeometric means test was performed using *goseq* v1.4 (Young et al. 2010) to identify significantly enriched GO terms.

## Acknowledgements

The authors thank I. Darolti for help with tissue sampling as well as W. van der Bijl, B. L. S. Furman, and B. Sandkam for providing bioinformatic support and insightful discussions during the preparation of this manuscript. This work was supported by the Canada 150 Research Chair Program and the European Research Council 680951) to J.E.M., who also gratefully acknowledges support from the Natural Science and Engineering Research Council of Canada. We thank Clara Lacy for the guppy artwork.

## Data Availability

Data will be available from the Genbank Sequence Read Archive following publication of this manuscript.

## References

Almeida P et al. 2020. Divergence and remarkable diversity of the Y chromosome in guppies. bioRxiv. 2020.07.13.200196. doi: 10.1101/2020.07.13.200196.

Anastasiadi D, Vandeputte M, Sánchez-Baizán N, Allal F, Piferrer F. 2018. Dynamic epimarks in sex-related genes predict gonad phenotype in the European sea bass, a fish with mixed genetic and environmental sex determination. Epigenetics. 13:988–1011. doi: 10.1080/15592294.2018.1529504.

Barlow DP, Bartolomei MS. 2014. Genomic imprinting in mammals. Cold Spring Harbor Perspectives in Biology. 6:a018382–a018382. doi: 10.1101/cshperspect.a018382.

Bolger AM, Lohse M, Usadel B. 2014. Trimmomatic: A flexible trimmer for Illumina sequence data. Bioinformatics. 30:2114–2120. doi: 10.1093/bioinformatics/btu170.

Champroux A, Cocquet J, Henry-Berger J, Drevet JR, Kocer A. 2018. A decade of exploring the mammalian sperm epigenome: Paternal epigenetic and transgenerational inheritance. Frontiers in Cell and Developmental Biology. 6. doi: 10.3389/fcell.2018.00050.

Chen S et al. 2014. Whole-genome sequence of a flatfish provides insights into ZW sex chromosome evolution and adaptation to a benthic lifestyle. Nature genetics. 46:253–60. doi: 10.1038/ng.2890.

Darolti I et al. 2019. Extreme heterogeneity in sex chromosome differentiation and dosage compensation in livebearers. Proceedings of the National Academy of Sciences. 116:19031–19036. doi: 10.1073/pnas.1905298116.

Darolti I, Wright AE, Mank JE. 2020. Guppy Y chromosome integrity maintained by incomplete recombination suppression. Genome Biology and Evolution. 12:965–977. doi: 10.1093/gbe/evaa099.

Deniz Ö, Frost JM, Branco MR. 2019. Regulation of transposable elements by DNA modifications. Nature Reviews Genetics. 20:417–431. doi: 10.1038/s41576-019-0106-6.

Furman BLS et al. 2020. Sex chromosome evolution: So many exceptions to the rules. Genome Biology and Evolution. 12:750–763. doi: 10.1093/gbe/evaa081.

Gorelick R. 2003. Evolution of dioecy and sex chromosomes via methylation driving Muller’s ratchet. Biological Journal of the Linnean Society. 80:353–368. doi: 10.1046/j.1095-8312.2003.00244.x.

Hammoud SS et al. 2009. Distinctive chromatin in human sperm packages genes for embryo development. Nature. 460:473–478. doi: 10.1038/nature08162.

Hata K, Okano M, Lei H, Li E. 2002. Dnmt3L cooperates with the Dnmt3 family of de novo DNA methyltransferases to establish maternal imprints in mice. Development. 129:1983–1993.

Heard E, Martienssen RA. 2014. Transgenerational epigenetic inheritance: myths and mechanisms. Cell. 157:95–109. doi: 10.1016/j.cell.2014.02.045.

Hernando-Herraez I et al. 2015. The interplay between DNA methylation and sequence divergence in recent human evolution. Nucleic Acids Research. 43:8204–8214. doi: 10.1093/nar/gkv693.

Iwase M et al. 2003. The amelogenin loci span an ancient pseudoautosomal boundary in diverse mammalian species. Proceedings of the National Academy of Sciences. 100:5258–5263. doi: 10.1073/pnas.0635848100.

Jaenisch R, Bird A. 2003. Epigenetic regulation of gene expression: how the genome integrates intrinsic and environmental signals. Nature Genetics. 33:245–254. doi: 10.1038/ng1089.

Jiang L et al. 2013. Sperm, but not oocyte, DNA methylome is inherited by zebrafish early embryos. Cell. 153:773–84. doi: 10.1016/j.cell.2013.04.041.

Jones PA. 2012. Functions of DNA methylation: islands, start sites, gene bodies and beyond. Nature Reviews Genetics. 13:484–492. doi: 10.1038/nrg3230.

Korthauer K, Chakraborty S, Benjamini Y, Irizarry RA. 2019. Detection and accurate false discovery rate control of differentially methylated regions from whole genome bisulfite sequencing. Biostatistics. 20:367–383. doi: 10.1093/biostatistics/kxy007.

Kotrschal A et al. 2013. The benefit of evolving a larger brain: big-brained guppies perform better in a cognitive task. Animal Behaviour. 86:e4–e6. doi: 10.1016/j.anbehav.2013.07.011.

Krueger F, Andrews SR. 2011. Bismark: a flexible aligner and methylation caller for Bisulfite-Seq applications. Bioinformatics (Oxford, England). 27:1571–2. doi: 10.1093/bioinformatics/btr167.

Li H et al. 2009. The Sequence Alignment/Map format and SAMtools. Bioinformatics. 25:2078–2079. doi: 10.1093/bioinformatics/btp352.

Liu J, Morgan M, Hutchison K, Calhoun VD. 2010. A study of the influence of sex on genome wide methylation Creighton, C, editor. PLoS ONE. 5:e10028. doi: 10.1371/journal.pone.0010028.

Long HK et al. 2013. Epigenetic conservation at gene regulatory elements revealed by nonmethylated DNA profiling in seven vertebrates. eLife. 2:1–19. doi: 10.7554/eLife.00348.

Lowdon RF, Jang HS, Wang T. 2016. Evolution of epigenetic regulation in vertebrate genomes. Trends in Genetics. 32:269–283. doi: 10.1016/j.tig.2016.03.001.

Luo Z et al. 2016. Regulation of the imprinted Dlk1-Dio3 locus by allele-specific enhancer activity. Genes & Development. 30:92–101. doi: 10.1101/gad.270413.115.

Mank JE. 2017. The transcriptional architecture of phenotypic dimorphism. Nature Ecology & Evolution. 1:0006. doi: 10.1038/s41559-016-0006.

Metzger DCH, Schulte PM. 2018. The DNA methylation landscape of stickleback reveals patterns of sex chromosome evolution and effects of environmental salinity. Genome Biology and Evolution. 10:775–785. doi: 10.1093/gbe/evy034.

Mirouze M et al. 2012. Loss of DNA methylation affects the recombination landscape in *Arabidopsis*. Proceedings of the National Academy of Sciences. 109:5880–5885. doi: 10.1073/pnas.1120841109.

Navarro-Martín L et al. 2011. DNA methylation of the gonadal aromatase (cyp19a) promoter is involved in temperature-dependent sex ratio shifts in the European sea bass. PLoS Genetics. 7:e1002447. doi: 10.1371/journal.pgen.1002447.

Nik-Zainal S et al. 2012. Mutational Processes Molding the Genomes of 21 Breast Cancers. Cell. 149:979–993. doi: 10.1016/j.cell.2012.04.024.

Ono R et al. 2006. Deletion of Peg10, an imprinted gene acquired from a retrotransposon, causes early embryonic lethality. Nature Genetics. 38:101–106. doi: 10.1038/ng1699.

Ortega-Recalde O, Day RC, Gemmell NJ, Hore TA. 2019. Zebrafish preserve global germline DNA methylation while sex-linked rDNA is amplified and demethylated during feminisation. Nature Communications. 10:3053. doi: 10.1038/s41467-019-10894-7.

Ortega-Recalde O, Hore TA. 2019. DNA methylation in the vertebrate germline: balancing memory and erasure. Essays in Biochemistry. 63:649–661. doi: 10.1042/EBC20190038.

Poeser Fred N, Kempkes M, Isbrücker IJH. 2005. Description of *Poecilia (Acanthophacelus) wingei* n. sp. from the Paría Peninsula, Venezuela, including notes on *Acanthophacelus* Eigenmann, 1907 and other subgenera of *Poecilia* Bloch and Schneider, 1801 (Teleostei, Cyprin. Contributions to Zoology. 74:97–115. doi: 10.1163/18759866-0740102007.

Pollux BJA, Meredith RW, Springer MS, Garland T, Reznick DN. 2014. The evolution of the placenta drives a shift in sexual selection in livebearing fish. Nature. 513:233–236. doi: 10.1038/nature13451.

Potok ME, Nix DA, Parnell TJ, Cairns BR. 2013. Reprogramming the maternal zebrafish genome after fertilization to match the paternal methylation pattern. Cell. 153:759–772. doi: 10.1016/j.cell.2013.04.030.

R Core Team. 2019. R: A language and environment for statistical computing. R Foundation for Statistical Computing, Vienna, Austria. https://www.r-project.org/.

Rabosky DL et al. 2018. An inverse latitudinal gradient in speciation rate for marine fishes. Nature. 559:392–395. doi: 10.1038/s41586-018-0273-1.

Rice WR. 1984. Sex chromosomes and the evolution of sexual dimorphism. Evolution. 38:735. doi: 10.2307/2408385.

Saldivar Lemus Y, Vielle-Calzada JP, Ritchie MG, Macías Garcia C. 2017. Asymmetric paternal effect on offspring size linked to parent-of-origin expression of an insulin-like growth factor. Ecology and Evolution. 7:4465–4474. doi: 10.1002/ece3.3025.

Schneider M et al. 2012. The innate immune sensor NLRC3 attenuates Toll-like receptor signaling via modification of the signaling adaptor TRAF6 and transcription factor NF-κB. Nature Immunology. 13:823–831. doi: 10.1038/ni.2378.

Sekita Y et al. 2008. Role of retrotransposon-derived imprinted gene, Rtl1, in the feto-maternal interface of mouse placenta. Nature Genetics. 40:243–248. doi: 10.1038/ng.2007.51.

Shao C et al. 2014. Epigenetic modification and inheritance in sexual reversal of fish. Genome Research. 24:604–615. doi: 10.1101/gr.162172.113.

Skvortsova K et al. 2019. Retention of paternal DNA methylome in the developing zebrafish germline. Nature Communications. 10:1–13. doi: 10.1038/s41467-019-10895-6.

Tena JJ et al. 2014. Comparative epigenomics in distantly related teleost species identifies conserved cis-regulatory nodes active during the vertebrate phylotypic period. Genome Research. 24:1075–1085. doi: 10.1101/gr.163915.113.

Tripathi N, Hoffmann M, Willing EM, et al. 2009. Genetic linkage map of the guppy, *Poecilia reticulata*, and quantitative trait loci analysis of male size and colour variation. Proceedings of the Royal Society B: Biological Sciences. 276:2195–2208. doi: 10.1098/rspb.2008.1930.

Tripathi N, Hoffmann M, Weigel D, Dreyer C. 2009. Linkage Analysis Reveals the Independent Origin of Poeciliid Sex Chromosomes and a Case of Atypical Sex Inheritance in the Guppy (*Poecilia reticulata*). Genetics. 182:365–374. doi: 10.1534/genetics.108.098541.

Volff JN. 2006. Turning junk into gold: Domestication of transposable elements and the creation of new genes in eukaryotes. BioEssays. 28:913–922. doi: 10.1002/bies.20452.

Wang X, Bhandari RK. 2019. DNA methylation dynamics during epigenetic reprogramming of medaka embryo. Epigenetics. 14:611–622. doi: 10.1080/15592294.2019.1605816.

Wright AE et al. 2017. Convergent recombination suppression suggests role of sexual selection in guppy sex chromosome formation. Nature Communications. 8:1–10. doi: 10.1038/ncomms14251.

Xu L et al. 2019. Dynamic evolutionary history and gene content of sex chromosomes across diverse songbirds. Nature Ecology and Evolution. 3:834–844. doi: 10.1038/s41559-019-0850-1.

Yokomine T, Hata K, Tsudzuki M, Sasaki H. 2006. Evolution of the vertebrate DNMT3 gene family: a possible link between existence of *DNMT3L* and genomic imprinting. Cytogenetic and Genome Research. 113:75–80. doi: 10.1159/000090817.

Young MD, Wakefield MJ, Smyth GK, Oshlack A. 2010. Gene ontology analysis for RNA-seq: accounting for selection bias. Genome biology. 11:R14. doi: 10.1186/gb-2010-11-2-r14.

Yu D et al. 2018. Silencing of retrotransposon-derived imprinted gene RTL1 is the main cause for postimplantational failures in mammalian cloning. Proceedings of the National Academy of Sciences. 115:E11071–E11080. doi: 10.1073/pnas.1814514115.

Zhou J et al. 2017. Tissue-specific DNA methylation is conserved across human, mouse, and rat, and driven by primary sequence conservation. BMC Genomics. 18:1–17. doi: 10.1186/s12864-017-4115-6.

