## Supplementary material for "Conserved sex-biased DNA methylation patterns target key developmental genes and non-recombining region of the guppy sex chromosome"

Table S1: Summary statistics for whole genome bisulfite sequencing and read mapping

| | | *P.reticulata* | | | | *P. wingei* | | | |
|---|---|---|---|---|---|---|---|---|---|
| | Sample # | Library Size | Post Filtering Library Size | Uniquely mapped reads | | Library Size | Post Filtering Library Size | Uniquely mapped reads | |
| Testis | 1 | 102,146,788 | 97,780,050 | 64,502,764 | 66.0% | 83,924,392 | 80,425,731 | 47,391,585 | 58.9% |
| | 2 | 91,681,567 | 87,827,553 | 58,887,484 | 67.0% | 82,088,986 | 78,670,949 | 45,248,193 | 57.5% |
| | 3 | 74,168,188 | 71,059,377 | 46,652,953 | 65.7% | 86,478,505 | 82,533,337 | 45,993,335 | 55.7% |
| | 4 | 94,988,723 | 90,825,462 | 61,000,924 | 67.2% | 88,797,874 | 84,623,636 | 49,954,138 | 59.0% |
| | 5 | 100,633,769 | 96,611,089 | 63,742,893 | 66.0% | 92,196,178 | 89,079,897 | 52,981,049 | 59.5% |
| Ovary | 1 | 74,844,076 | 71,669,383 | 48,837,271 | 68.1% | 72,866,094 | 69,843,703 | 43,131,639 | 61.8% |
| | 2 | 87,268,129 | 83,695,821 | 57,053,369 | 68.2% | 84,500,758 | 81,199,624 | 51,076,513 | 62.9% |
| | 3 | 73,594,086 | 70,485,208 | 47,328,220 | 67.1% | 92,805,423 | 88,974,067 | 54,754,300 | 61.5% |
| | 4 | 83,612,641 | 80,145,014 | 54,782,950 | 68.4% | 66,648,233 | 64,053,407 | 39,672,327 | 61.9% |
| | 5 | 89,953,963 | 86,511,974 | 58,675,083 | 67.8% | 102,839,337 | 99,120,404 | 61,107,723 | 61.6% |
| Male Muscle | 1 | 62,321,415 | 59,654,426 | 39,776,437 | 66.7% | 75,044,531 | 72,123,554 | 46,042,790 | 63.8% |
| | 2 | 66,564,898 | 64,132,803 | 44,235,111 | 69.0% | 85,124,504 | 81,877,072 | 52,394,092 | 64.0% |
| | 3 | 74,948,234 | 71,764,085 | 48,316,501 | 67.3% | 69,043,710 | 66,543,802 | 42,133,846 | 63.3% |
| | 4 | 59,620,621 | 57,101,772 | 38,480,217 | 67.4% | 68,628,502 | 65,866,966 | 41,505,222 | 63.0% |
| | 5 | 64,108,375 | 61,655,494 | 41,259,146 | 66.9% | 96,504,103 | 93,111,046 | 58,121,489 | 62.4% |
| Female Muscle | 1 | 69,644,727 | 66,819,753 | 46,339,825 | 69.4% | 68,963,847 | 66,447,226 | 42,971,443 | 64.7% |
| | 2 | 84,584,807 | 81,003,172 | 54,120,925 | 66.8% | 60,334,361 | 58,127,279 | 37,013,320 | 63.7% |
| | 3 | 68,732,610 | 66,062,287 | 44,664,427 | 67.6% | 74,414,116 | 71,565,106 | 45,141,706 | 63.1% |
| | 4 | 59,280,474 | 56,649,663 | 37,699,906 | 66.5% | 76,879,825 | 74,273,937 | 48,195,801 | 64.9% |
| | 5 | 79,246,203 | 76,193,327 | 50,883,958 | 66.8% | 80,124,653 | 77,370,675 | 49,464,012 | 63.9% |

Table S2: 100kb genomic windows enriched for hypomethylated DMCs

| Chromosome | start | # CpGs | # hypomethylated | # hypermethylated | p-value |
|---|---|---|---|---|---|
| 1 | 14,700,000 | 460 | 169 (37%) | 53 | 1.06E-132 |
| 8 | 1,700,000 | 441 | 161 (37%) | 13 | 5.84E-126 |
| 8 | 1,600,000 | 346 | 123 (36%) | 8 | 9.53E-95 |
| 11 | 28,100,000 | 147 | 61 (41%) | 12 | 1.27E-51 |
| 18 | 12,600,000 | 242 | 72 (30%) | 8 | 7.00E-49 |
| 12 | 20,700,000 | 71 | 45 (63%) | 2 | 8.00E-49 |
| 19 | 9,000,000 | 236 | 69 (29%) | 12 | 3.85E-46 |
| 11 | 9,200,000 | 196 | 57 (29%) | 6 | 2.59E-38 |
| 4 | 3,400,000 | 92 | 41 (45%) | 3 | 9.50E-36 |
| 2 | 15,000,000 | 117 | 42 (36%) | 5 | 9.61E-32 |
| 5 | 24,700,000 | 75 | 30 (40%) | 13 | 1.90E-24 |
| 5 | 900,000 | 41 | 23 (56%) | 0 | 4.45E-23 |
| 10 | 32,400,000 | 86 | 29 (34%) | 1 | 1.98E-21 |
| 9 | 32,700,000 | 43 | 22 (51%) | 1 | 1.12E-20 |
| 7 | 30,200,000 | 52 | 22 (42%) | 4 | 6.87E-19 |
| 15 | 20,700,000 | 39 | 19 (49%) | 7 | 1.45E-17 |
| 6 | 31,000,000 | 68 | 23 (34%) | 0 | 5.81E-17 |
| 11 | 28,600,000 | 17 | 13 (76%) | 0 | 4.46E-16 |
| 19 | 1,500,000 | 46 | 19 (41%) | 0 | 4.47E-16 |
| 10 | 31,400,000 | 72 | 22 (31%) | 8 | 2.35E-15 |
| 15 | 29,800,000 | 70 | 21 (30%) | 3 | 2.38E-14 |
| 4 | 4,600,000 | 42 | 16 (38%) | 1 | 8.74E-13 |
| 1 | 34,000,000 | 41 | 14 (34%) | 1 | 1.44E-10 |

Table S3: 100kb genomic windows enriched for hypermethylated DMCs

| Chromosome | start | # CpGs | # hypomethylated | # hypermethylated | p-value |
|---|---|---|---|---|---|
| 8 | 13,700,000 | 352 | 0 | 110 (31%) | 1.74E-19 |
| 5 | 12,800,000 | 249 | 6 | 88 (35%) | 2.75E-19 |
| 5 | 7,700,000 | 318 | 12 | 102 (32%) | 3.67E-19 |
| 4 | 27,700,000 | 265 | 0 | 91 (34%) | 5.07E-19 |
| 2 | 37,300,000 | 246 | 9 | 87 (35%) | 6.01E-19 |
| 22 | 23,300,000 | 209 | 0 | 78 (37%) | 8.88E-19 |
| 14 | 10,800,000 | 261 | 0 | 88 (34%) | 8.15E-18 |
| 21 | 9,200,000 | 335 | 14 | 102 (30%) | 3.84E-17 |
| 15 | 17,600,000 | 332 | 1 | 100 (30%) | 2.37E-16 |
| 1 | 20,000,000 | 357 | 0 | 104 (29%) | 7.66E-16 |
| 8 | 12,100,000 | 279 | 1 | 87 (31%) | 1.74E-15 |
| 10 | 16,800,000 | 262 | 25 | 84 (32%) | 2.22E-15 |
| 8 | 14,600,000 | 318 | 3 | 94 (30%) | 3.00E-15 |
| 2 | 31,100,000 | 326 | 1 | 96 (29%) | 4.57E-15 |
| 21 | 9,800,000 | 302 | 0 | 90 (30%) | 1.09E-14 |
| 14 | 2,100,000 | 258 | 0 | 81 (31%) | 1.44E-14 |
| 14 | 11,000,000 | 209 | 0 | 70 (33%) | 2.61E-14 |
| 8 | 26,000,000 | 201 | 4 | 68 (34%) | 2.84E-14 |
| 3 | 26,00,000 | 293 | 0 | 88 (30%) | 3.82E-14 |
| 15 | 25,400,000 | 272 | 25 | 83 (31%) | 4.27E-14 |
| 16 | 9,000,000 | 280 | 3 | 85 (30%) | 5.78E-14 |
| 3 | 25,100,000 | 241 | 0 | 76 (32%) | 1.16E-13 |
| 17 | 26,400,000 | 298 | 1 | 87 (29%) | 3.24E-13 |
| 2 | 28,900,000 | 278 | 6 | 82 (29%) | 5.52E-13 |
| 12 | 200,000 | 277 | 4 | 82 (30%) | 7.56E-13 |
| 10 | 28,900,000 | 239 | 0 | 74 (31%) | 7.61E-13 |
| 7 | 1,900,000 | 252 | 0 | 77 (31%) | 9.21E-13 |
| 7 | 25,900,000 | 222 | 5 | 70 (32%) | 1.40E-12 |
| 22 | 23,400,000 | 205 | 4 | 66 (32%) | 1.40E-12 |
| 13 | 20,900,000 | 269 | 0 | 79 (29%) | 2.44E-12 |
| 14 | 10,400,000 | 205 | 0 | 65 (32%) | 4.13E-12 |
| 10 | 28,800,000 | 239 | 8 | 72 (30%) | 5.50E-12 |
| 17 | 26,500,000 | 228 | 12 | 70 (31%) | 6.06E-12 |
| 1 | 700,000 | 254 | 17 | 75 (30%) | 8.26E-12 |
| 8 | 22,400,000 | 195 | 2 | 62 (32%) | 8.81E-12 |
| 5 | 9,300,000 | 224 | 0 | 68 (30%) | 1.39E-11 |
| 5 | 14,500,000 | 233 | 0 | 68 (29%) | 6.21E-11 |
| 12 | 6,600,000 | 222 | 0 | 66 (30%) | 1.41E-10 |
| 14 | 900,000 | 207 | 3 | 62 (30%) | 2.01E-10 |
| 10 | 2,500,000 | 207 | 8 | 62 (30%) | 2.32E-10 |
| 2 | 28,100,000 | 191 | 0 | 59 (31%) | 3.21E-10 |
| 17 | 28,100,000 | 173 | 17 | 53 (31%) | 4.96E-09 |
| 8 | 12,500,000 | 169 | 0 | 51 (30%) | 5.64E-09 |

Table S4: Significantly enriched GO terms for genes located near hypomethylated DMRs

| GO: ID | Over represented p-value | Under represented p-value | # DiffMeth Genes | number of genes in category | GO term |
|---|---|---|---|---|---|
| GO:0007156 | 4.12E-08 | 0.999999993 | 18 | 133 | homophilic cell adhesion via plasma membrane adhesion molecules |
| GO:0070588 | 0.000168783 | 0.999974578 | 8 | 56 | calcium ion transmembrane transport |
| GO:0007155 | 0.000817352 | 0.999705828 | 17 | 250 | cell adhesion |
| GO:0007017 | 0.000848378 | 0.999883654 | 6 | 40 | microtubule-based process |
| GO:0009966 | 0.002871483 | 0.999595621 | 5 | 35 | regulation of signal transduction |
| GO:0039021 | 0.003139939 | 0.999821115 | 3 | 11 | pronephric glomerulus development |
| GO:0048514 | 0.006643712 | 0.999139858 | 4 | 27 | blood vessel morphogenesis |
| GO:0007165 | 0.012128132 | 0.993560171 | 23 | 487 | signal transduction |
| GO:0098609 | 0.013262484 | 0.997232631 | 5 | 50 | cell-cell adhesion |
| GO:0006471 | 0.013404439 | 0.998583385 | 3 | 18 | protein ADP-ribosylation |
| GO:0042127 | 0.01356024 | 0.997800938 | 4 | 33 | regulation of cell proliferation |
| GO:0015986 | 0.015588795 | 0.99824546 | 3 | 19 | ATP synthesis coupled proton transport |
| GO:0007409 | 0.016601551 | 0.997128517 | 4 | 35 | axonogenesis |
| GO:0007519 | 0.016601551 | 0.997128517 | 4 | 35 | skeletal muscle tissue development |
| GO:0060059 | 0.017961313 | 0.997855469 | 3 | 20 | embryonic retina morphogenesis in camera-type eye |
| GO:0051056 | 0.020523127 | 0.997409571 | 3 | 21 | regulation of small GTPase mediated signal transduction |
| GO:0015074 | 0.023132194 | 0.993445835 | 6 | 78 | DNA integration |
| GO:0035556 | 0.025483238 | 0.987881039 | 14 | 275 | intracellular signal transduction |
| GO:0001944 | 0.025958512 | 0.994815741 | 4 | 40 | vasculature development |
| GO:0006811 | 0.027132258 | 0.986614181 | 15 | 304 | ion transport |
| GO:0060047 | 0.030429662 | 0.993600182 | 4 | 42 | heart contraction |
| GO:0006355 | 0.033033876 | 0.978234609 | 37 | 953 | regulation of transcription, DNA-templated |
| GO:0006816 | 0.043478981 | 0.989714195 | 4 | 47 | calcium ion transport |
| GO:0006351 | 0.045685037 | 0.986672658 | 5 | 69 | transcription, DNA-templated |

Table S5: Significantly enriched GO term for genes located near hypermethylated DMRs

| GO: ID | Over represented p-value | Under represented p-value | # DiffMeth Genes | number of genes in category | GO term |
|---|---|---|---|---|---|
| GO:0006468 | 0.048210176 | 0.985702993 | 5 | 631 | protein phosphorylation |

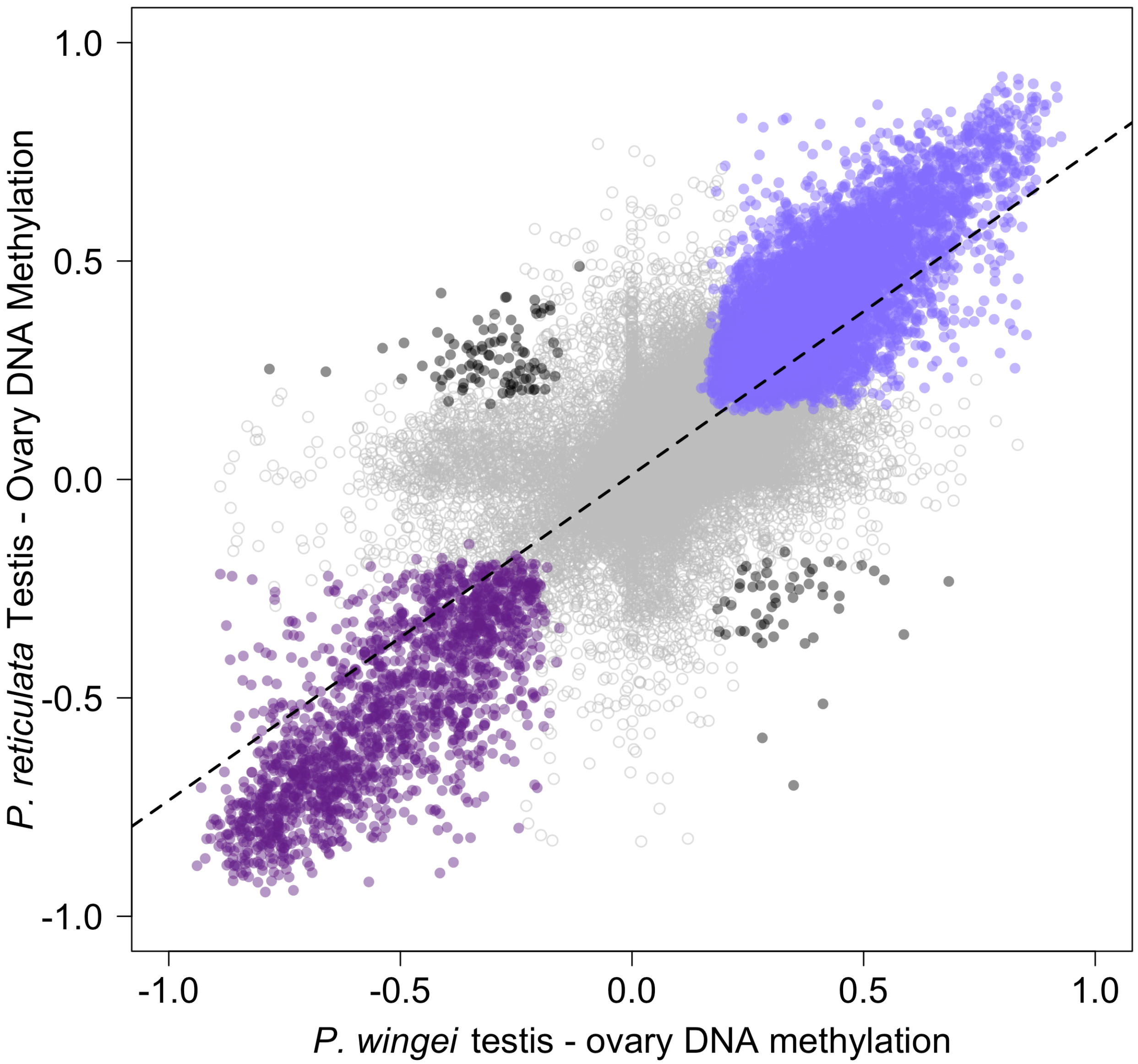


Figure S1: Correlation of differentially methylated CpG loci (DMCs) between testis and ovary for *P. reticulata* and *P. wingei*. Colours correspond to DMCs that are in both species that are either hypomethylated (purple), hypermethylated (blue), or divergent between species (black). Open grey circles represent loci present in both species but are not significant DMCs. Dashed line is the regression line ($R^2 = 0.617$, $p < 0.001$).

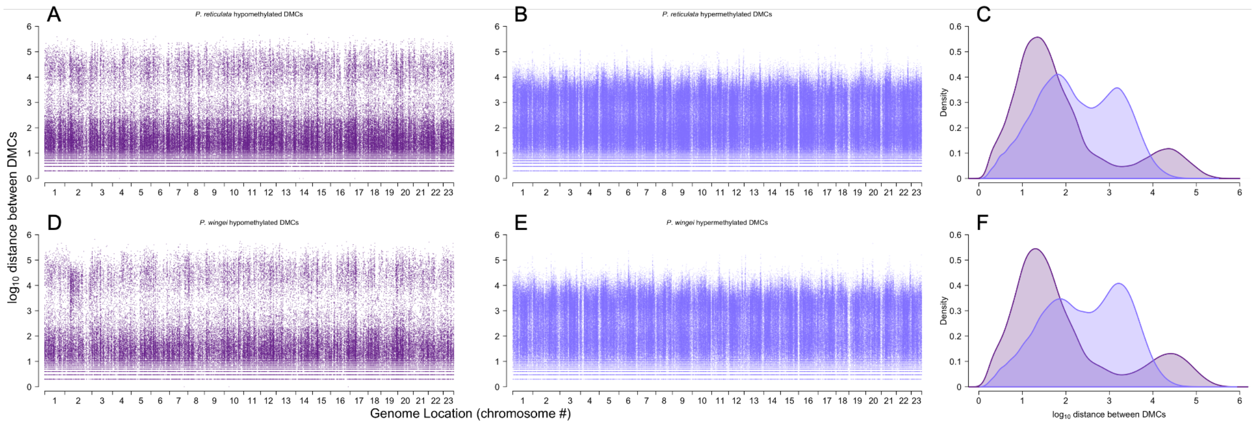


Figure S2: Analysis of the $\log_{10}$ genomic distance between hypomethylated (A & D, purple) or hypermethylated (B & E, blue) differentially methylated CpG loci for *Poecilia reticulata* (A & B) and *Poecilia wingei* (D & E) between testis and ovary. (C & F) Density distribution of the $\log_{10}$ distance between DMCs for *P. reticulata* (C) and *P. wingei* (F).

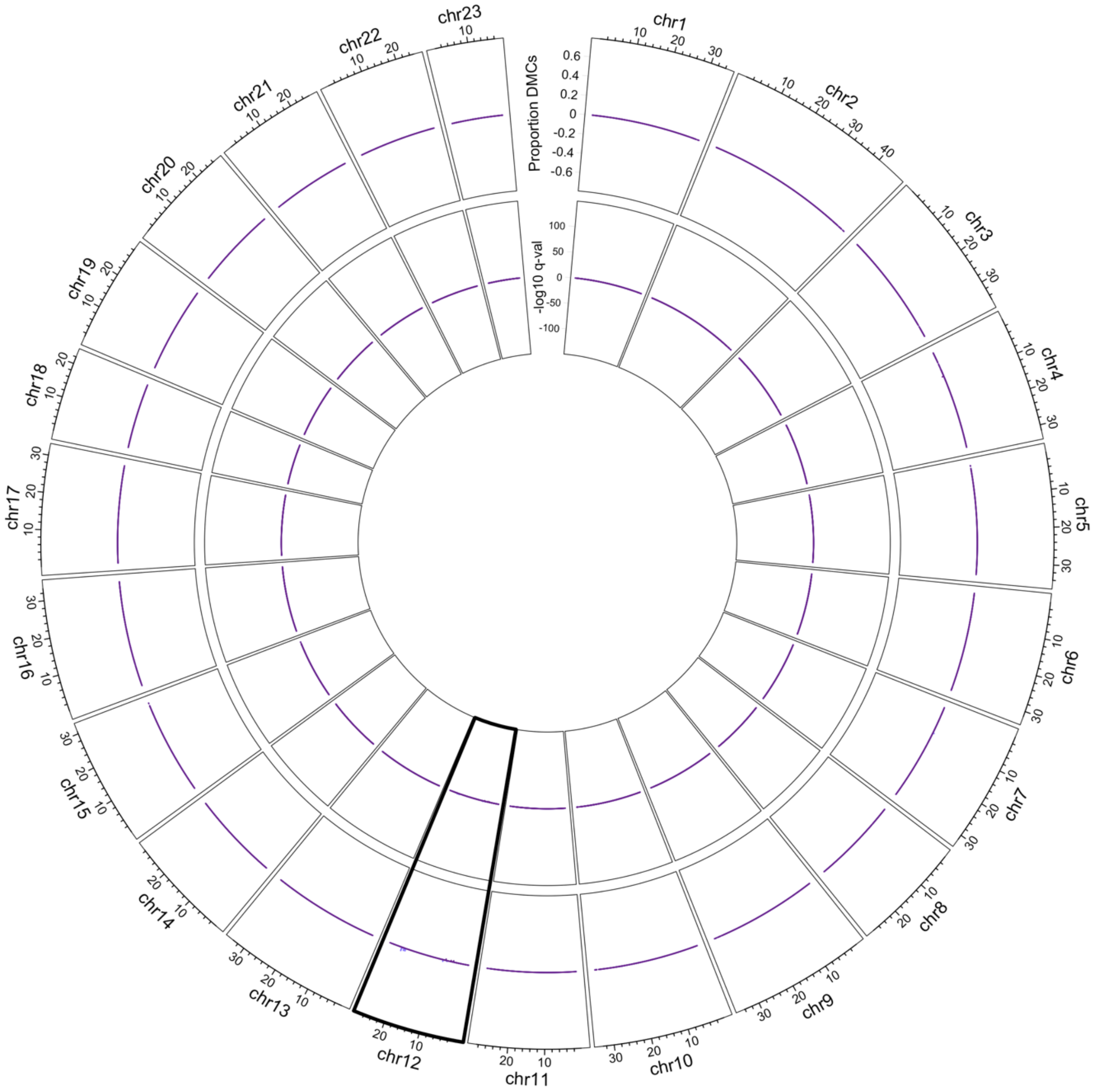


Figure S3: Circos plot of differential methylation between male and female muscle tissue for all 23 guppy chromosomes. The outer ring depicts the proportion of conserved DMCs between *Poecilia reticulata* and *Poecilia wingei* that are either hypermethylated (blue) or hypomethylated (purple) in male relative to female muscle tissue. Values on the y-axis represent the total number of hypermethylated or hypomethylated DMCs relative to the number of sequenced CpG loci within a 100 kb window. The inner ring is the -$\log_{10}$ FDR corrected q-value from a binomial test of the number of hypermethylated or hypomethylated DMCs in a 100 kb window. These data show few sex-specific differences in DNA methylation patterns between males and female muscle tissue, and little to no effect of genetic variation between the sample DNA and the reference genome on genomic DNA methylation patterns.

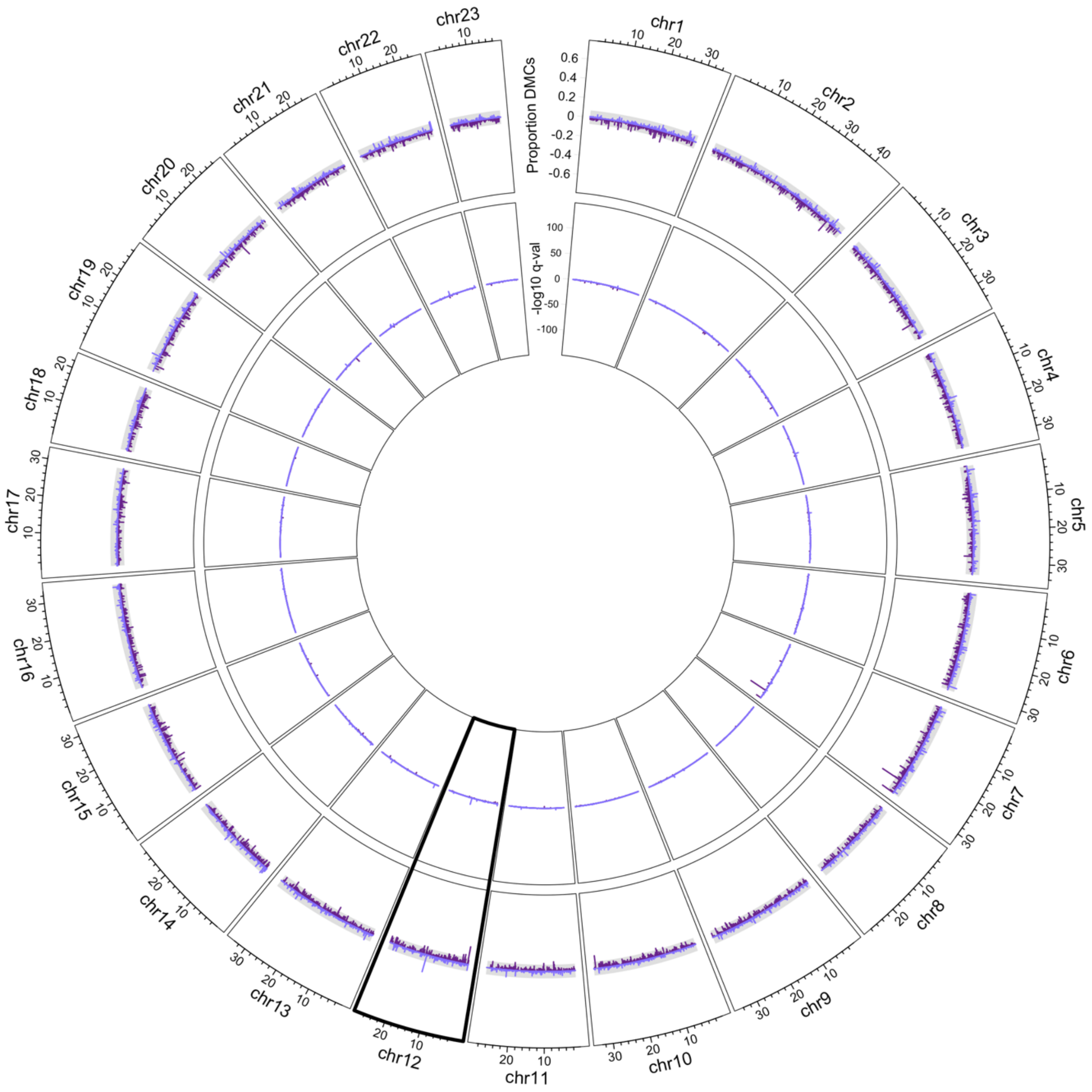


Figure S4: Circos plot of differential methylation between male muscle and ovary tissue for all 23 guppy chromosomes. The outer ring depicts the proportion of conserved DMCs between *Poecilia reticulata* and *Poecilia wingei* that are either hypermethylated (blue) or hypomethylated (purple) in male muscle relative to ovary tissue. Values on the y-axis represent the total number of hypermethylated or hypomethylated DMCs relative to the number of sequenced CpG loci within a 100 kb window. The inner circle is the -$\log_{10}$ FDR corrected q-value from a binomial test of the number of hypermethylated or hypomethylated DMCs in a 100 kb window.

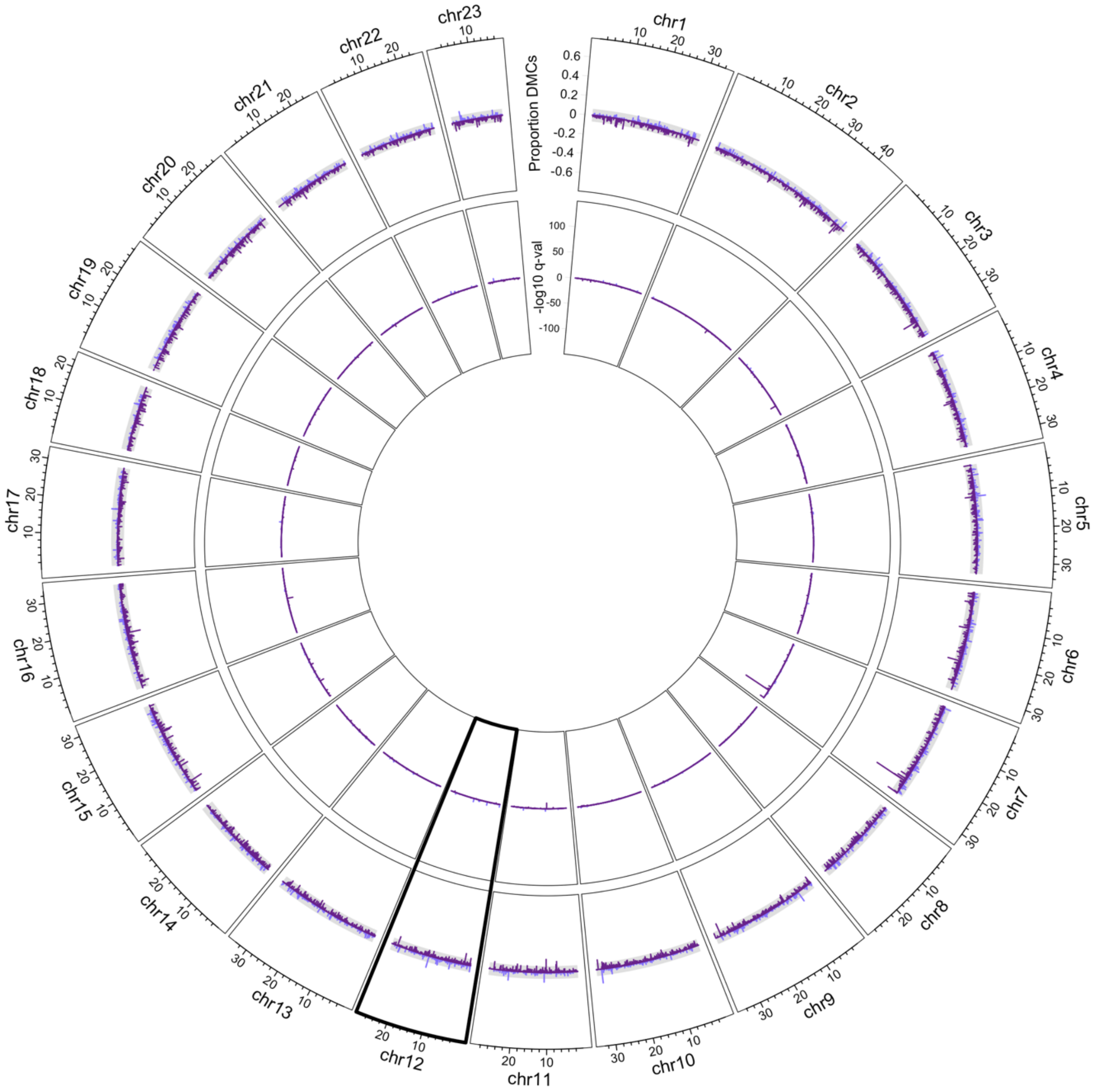


Figure S5: Circos plot of differential methylation between female muscle and ovary tissue for all 23 guppy chromosomes. The outer ring depicts the proportion of conserved DMCs between *Poecilia reticulata* and *Poecilia wingei* that are either hypermethylated (blue) or hypomethylated (purple) in female muscle relative to ovary tissue. Values on the y-axis represent the total number of hypermethylated or hypomethylated DMCs relative to the number of sequenced CpG loci within a 100 kb window. The inner circle is the -$\log_{10}$ FDR corrected q-value from a bionomial test of the number of hypermethylated or hypomethylated DMCs in a 100 kb window.

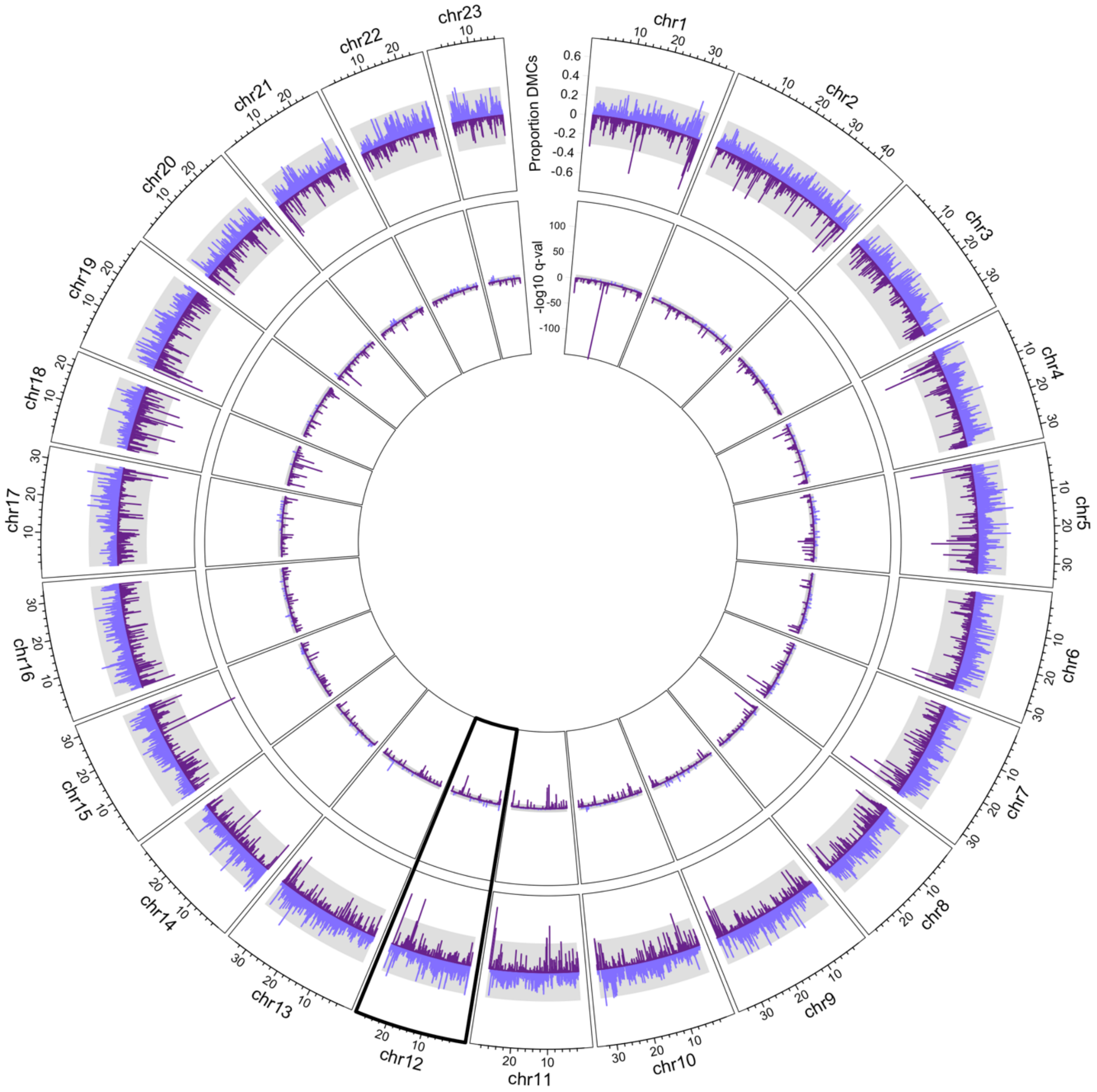


Figure S6: Circos plot of differential methylation between testis and male muscle for all 23 guppy chromosomes. The outer ring depicts the proportion of conserved DMCs between *Poecilia reticulata* and *Poecilia wingei* that are either hypermethylated (blue) or hypomethylated (purple) in testis relative to muscle tissue. Values on the y-axis represent the total number of hypermethylated or hypomethylated DMCs relative to the number of sequenced CpG loci within a 100 kb window. The inner circle is the -$\log_{10}$ FDR corrected q-value from a bionomial test of the number of hypermethylated or hypomethylated DMCs in a 100 kb window.

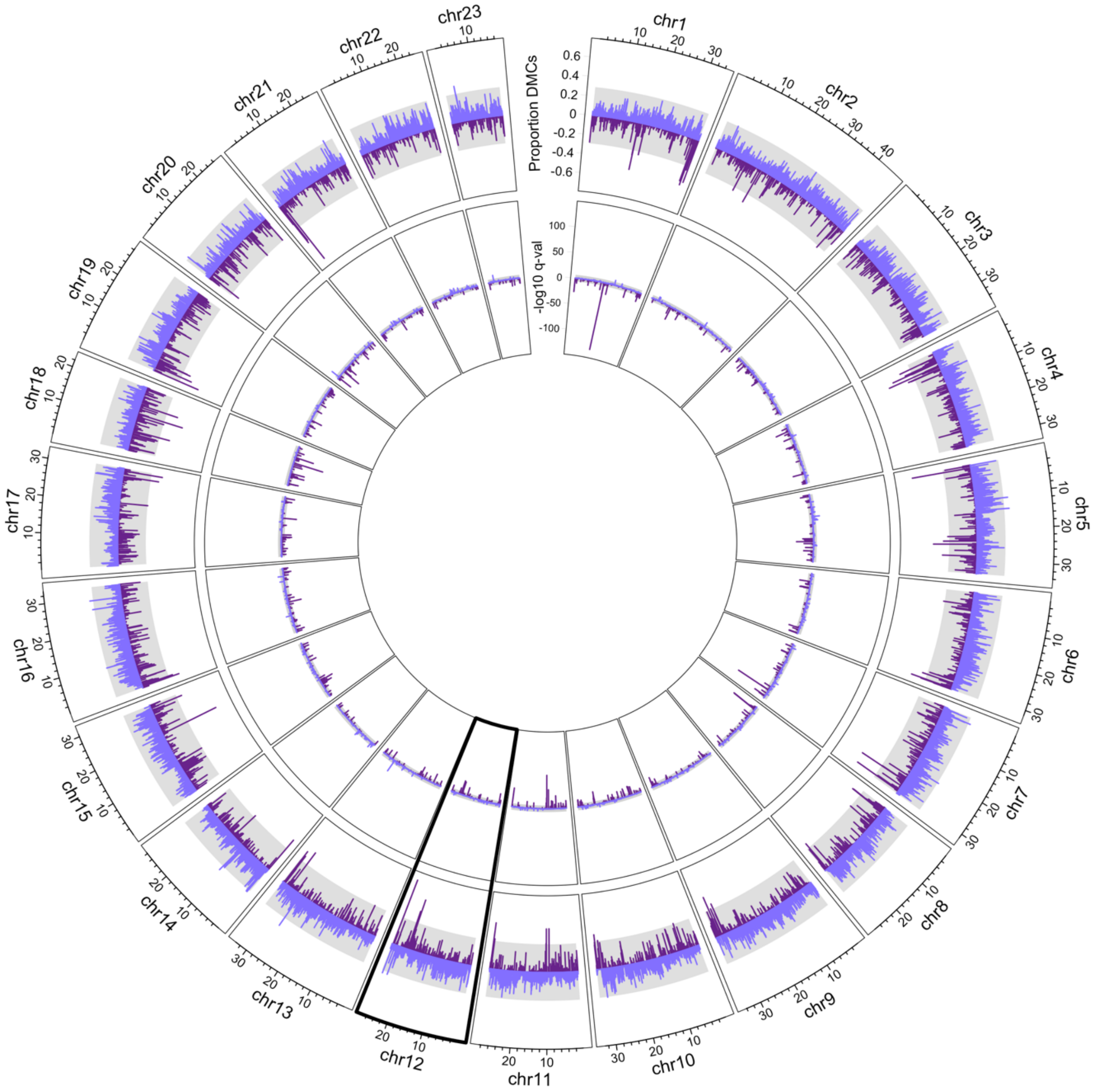


Figure S7: Circos plot of differential methylation between testis and female muscle for all 23 guppy chromosomes. The outer ring depicts the proportion of conserved DMCs between *Poecilia reticulata* and *Poecilia wingei* that are either hypermethylated (blue) or hypomethylated (purple) in testis relative to muscle tissue. Values on the y-axis represent the total number of hypermethylated or hypomethylated DMCs relative to the number of sequenced CpG loci within a 100 kb window. The inner circle is the -$\log_{10}$ FDR corrected q-value from a bionomial test of the number of hypermethylated or hypomethylated DMCs in a 100 kb window.

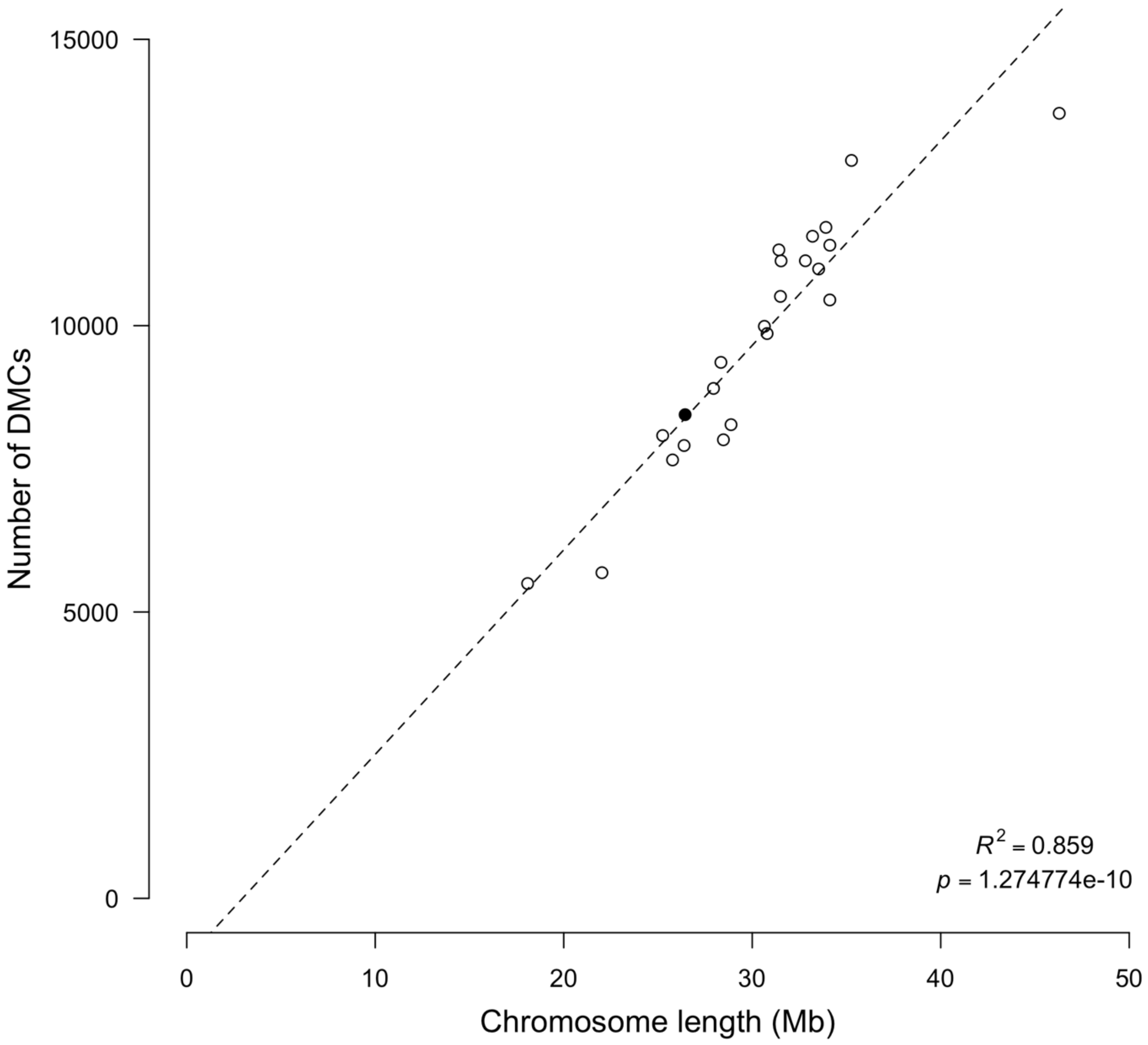


Figure S8:
Correlation between the number of significantly differentially methylated CpG sites on a chromosome and chromosome length. Open circles represent autosomes, filled circle is the sex chromosome (Chromosome 12).

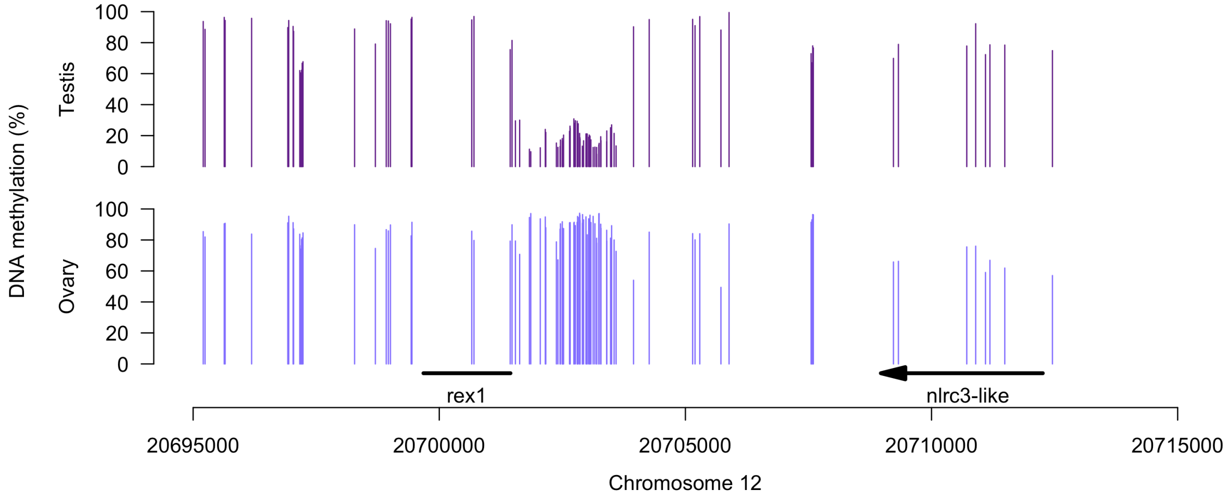


Figure S9: Percent methylation of individual CpG loci around the putative rex1 transposable element and the nlrc3-like gene in testis (upper panel, purple) and ovary (lower panel, blue) on Chromosome 12.

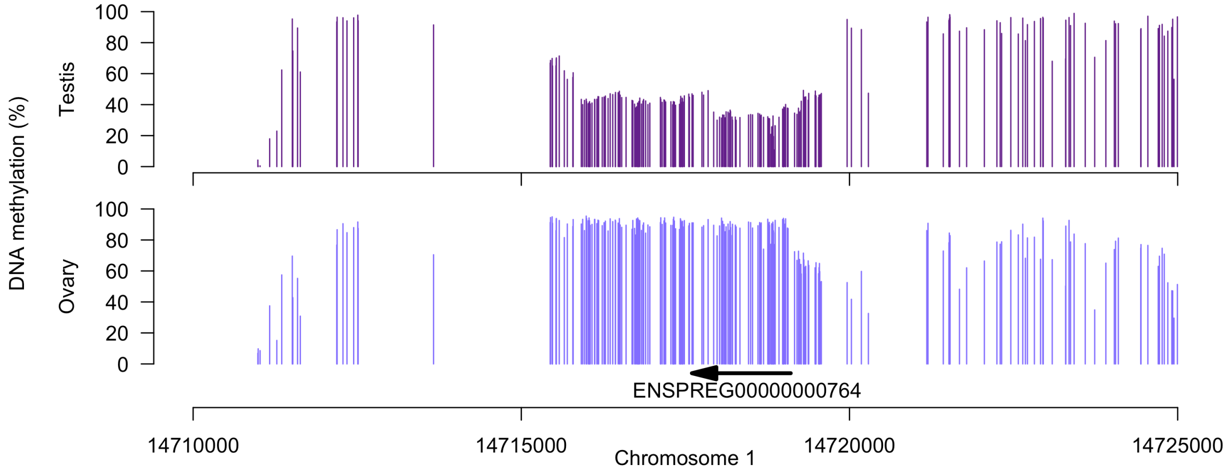


Figure S10: Percent methylation of individual CpG loci around the ENSPREG00000000764 gene in testis (upper panel, purple) and ovary (lower panel, blue) on Chromosome 1.
